# Decoding of the ubiquitin code for clearance of colliding ribosomes by the RQT complex

**DOI:** 10.1101/2022.09.12.507701

**Authors:** Yoshitaka Matsuo, Takayuki Uchihashi, Toshifumi Inada

## Abstract

The collision sensor Hel2 specifically recognizes colliding ribosomes and ubiquitinates the ribosomal protein uS10, leading to noncanonical subunit dissociation by the ribosome-associated quality control trigger (RQT) complex. Although uS10 ubiquitination is essential for rescuing stalled ribosomes, its function and recognition steps are not fully understood. Here, we showed that the RQT complex components Cue3 and Rqt4 interacted with the K63-linked ubiquitin chain and accelerated the recruitment of the RQT complex to the ubiquitinated colliding ribosome. The CUE domain of Cue3 and the N-terminal domain of Rqt4 bound independently to the K63-linked ubiquitin chain. Their deletion abolished ribosomal dissociation mediated by the RQT complex. High-speed atomic force microscopy (HS-AFM) reveals that the intrinsically disordered regions of Rqt4 enabled the expansion of the searchable area for interaction with the ubiquitin chain. These findings provide mechanistic insight into the decoding of the ubiquitin code for clearance of colliding ribosomes by the RQT complex.

## Introduction

Persistent translation arrest results in ribosome collision, which is considered a hallmark of translational stress^1–7^. Several sensor proteins recognize ribosomal collision and induce different cellular responses, including the ribosome-associated quality control (RQC) pathway^8–16^, no-go decay (N)^17–19^, the integrated stress response (ISR)^20–25^, and the ribotoxic stress response (RSR)^21,26–28^. The RQC pathway is the rescue system for stalled ribosomes, thereby contributing to the clearance of collided ribosomes to decrease the severity of translational stress. Defects in the RQC system lead to the enhancement of other translational stress responses including ISR and RSR^22,26,29^.

In the RQC pathway, the E3 ligase Hel2 (ZNF598 in mammals) recognizes the colliding ribosome and ubiquitinates the ribosomal protein uSl0 as a marker of aberrant translation^9,11,14^. The ubiquitinated leading stalled ribosome is disassembled by the ribosome quality control triggering (RQT) complex, which is composed of the RNA helicase Slhl (RQT2) (ASCC3 in mammals), the ubiquitin-binding protein Cue3 (RQT3) (ASCC2 in mammals), and the zinc-finger protein Rqt4 (TRIP4 in mammals), thus enabling the following ribosomes to resume translation^10,12^. After ribosomal splitting, the incomplete nascent chain is retained in the disassembled 60S subunit and is ubiquitinated by the E3 ligase Ltnl, followed by proteasomal degradation^6,30–35^. The ubiquitination of uSl0 is essential to activate the RQC pathway; however, its function and recognition steps are not fully understood. Although the coupling of ubiquitin conjugation to ER degradation (CUE) domain of Cue3, which is known as the ubiquitin-binding domain, directly binds to ubiquitin, deletion of Cue3 does not completely suppress RQC activity^11^, suggesting that an additional factor recognizes ubiquitinated uSl0 to assist in the splitting event.

In this study, we showed that Cue3 and Rqt4 of the RQT complex interact with the K63-linked ubiquitin chain and facilitate the recruitment of the RQT complex to ubiquitinated colliding ribosomes. The CUE domain of Cue3 and the N-terminal domain of Rqt4 bound to the K63-linked ubiquitin chain independently of each other. A deletion mutant of the two domains in the RQT complex lost the ability to bind to the K63-linked ubiquitin chain as well as recruitment to ubiquitinated colliding ribosomes, resulting in the loss of ribosome splitting activity. The dynamics of the RQT complex were visualized using high-speed atomic force microscopy (HS-AFM), which showed that the intrinsically disordered regions (IDRs) of Rqt4 enabled the expansion of the searchable area for interaction with the ubiquitin chain. These findings led us to propose a model in which the RQT complex binds to the K63-linked ubiquitin chain via two arms composed of Cue3 and Rqt4, and recruits itself to the ubiquitinated colliding ribosome, followed by ribosome splitting mediated by the helicase activity of Slhl.

## Results

### In vitro reconstitution of colliding ribosomes conjugated to K63-linked ubiquitin chains

To gain mechanistic insight into the decoding of the ubiquitin code of colliding ribosomes, we used a previously established *in vitro* system to reconstitute the ubiquitinated colliding ribosome using an RQC-inducible arrest sequence in the *SDD1* mRNA^10^. The cell-free *in vitro* translation extract was generated in a 3’-to-5’ mRNA decay-defective and uS10-detectable yeast strain, (*ski2t*Δ, *uS10-3HA*) to obtain a high amount of ribosome-nascent chain complexes (RNCs) and detect the ubiquitination of uS10 **(Fig. 1a)**.

**Fig. 1.**
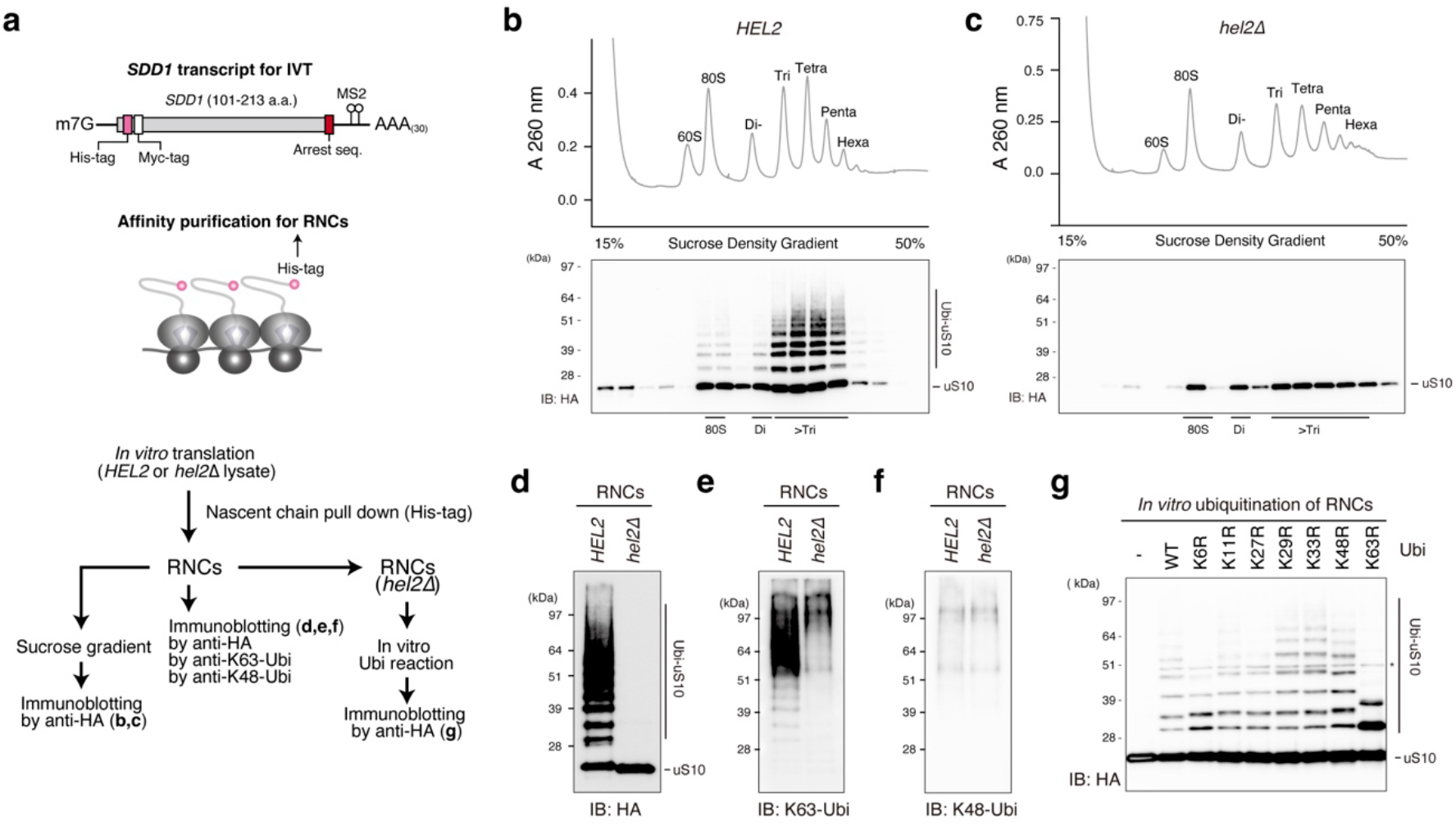
*In vitro* reconstitution of the ubiquitinated colliding ribosomes. **(a)** Schematic drawing of the *SDD1* model mRNA used in the *in vitro* translation (TVT) assay (Top). Schematic of the *in vitro* experiments (bottom). **(b, c)** The purified RNCs in the TVT reaction using *HEL2*-containing or *hel2*-knockout (*hel2*Δ) TVT extract were separated by sucrose density gradient centrifugation and detected by UV absorbance at a wavelength of 260 nm. HA-tagged uS10 in each fraction was detected by immunoblotting using an anti-HA antibody. **(d-f)** Immunoblotting of purified RNCs using anti-HA (**d**), K63-linkage (**e**), and K48-linkage-specific anti-ubiquitin antibodies (**f**). **(g)** *In vitro* ubiquitination assay. Purified RNCs derived from the *hel2*-knockout (*hel2*Δ) TVT reaction mixed with Hel2 (E3), Uba1 (E1), Ubc4 (E2), ATP, and ubiquitin or the indicated ubiquitin mutants. After the reaction, HA-tagged uS10 was detected by immunoblotting using an anti-HA antibody.

After *in vitro* translation, RNCs were affinity-purified using the N-terminal His-tag of the nascent chain and separated by sucrose density gradient centrifugation **(Fig. 1a)**. This clearly showed that the multimeric ribosomes were highly enriched in the purified RNCs **(Fig. 1b)** and that uS10 in heavy fractions more than disome were efficiently ubiquitinated during the *in vitro* translation **(Fig. 1b)**, indicating that the purified translating ribosomes could be collided and ubiquitinated by Hel2 as previously reported^10^.

Next, we prepared the *in vitro* translation extract from the *hel2* knockout yeast strain (*ski2*Δ, *uS10-3HA, hel2*Δ) and performed *in vitro* translation to obtain the nonubiquitinated colliding ribosomes. The purified RNCs contained many colliding ribosomes as well as *HEL2*-containing *in vitro* translation system, whereas ubiquitination of uS10 was abolished in the *hel2*-knockout *in vitro* translation system **(Fig. 1c, d)**.

To investigate the linkage type in the ubiquitin chains on uS10, ubiquitinated (*HEL2*) and nonubiquitinated (*hel2*Δ) colliding ribosomes were analyzed by immunoblotting using K63- or K48-linkage-specific ubiquitin antibodies. The K63-linked, but not K48-linked, ubiquitin chains were highly enriched in the ubiquitinated colliding ribosomes (*HEL2*) **(Fig. 1e, f)**. The bands of K63-linked ubiquitin chains were identical to the bands of ubiquitinated uS10 detected by the anti-HA antibody **(Fig. 1d, e)**. By contrast, these bands were not observed in the colliding ribosomes obtained from the *hel2*-knockout *in vitro* translation system **(Fig. 1c, d)**, indicating that uS10 in the colliding ribosomes was specifically modified with K63-linked ubiquitin chains by Hel2. The polyubiquitin linkage type in uS10 was confirmed using an *in vitro* ubiquitination assay with nonubiquitinated colliding ribosomes prepared with the *hel2*-knockout *in vitro* translation system, Hel2 (E3), Uba1 (E1), Ubc4 (E2), ATP, and various ubiquitin mutants with mutations at each polyubiquitination linkage site. The K63R mutation blocked the polyubiquitination of uS10, and only single or tandem monoubiquitination of two ubiquitination sites was observed on uS10 **(Fig. 1g)**. These results indicate that the colliding ribosomes were modified with K63-linked ubiquitin chains on uS10 by Hel2 *in vitro*.

### Cue3 and Rqt4 bind to the K63-linked ubiquitin chain

We previously showed that ubiquitinated uS10 is recognized by Cue3 in the RQT complex^11^. However, the direct interaction between the RQT complex and the polyubiquitin chain and its linkage type specificity remain to be clarified. A single deletion mutant of Cue3 or Rqt4 shows partial reduction of RQC activity, whereas the double deletion mutant shows complete loss of RQC function^11^. These results let us hypothesize that Cue3 and Rqt4 could have overlapping functions, namely, they both recognize the ubiquitin chain in the colliding ribosomes.

To investigate the interaction between the RQT complex and the polyubiquitin chain, we first determined whether it is possible to purify the RQT complex lacking Cue3 or Rqt4. For the immunoprecipitation and immunoblotting experiments, we constructed the C-terminal Flag-TEV-protein A-tagged Slh1, and the C-terminal Flag-tagged Cue3 and Rqt4 **(Fig. 2a)**. Several combinations of these RQT factors were coexpressed in yeast cells, and the complex was affinity-purified via the Slh1-Flag-TEV-protein A using IgG magnetic beads **(Fig. 2b)**. The Slh1/Cue3/Rqt4 complex was purified as a heterotrimer complex^11^ **(Fig. 2c)**, and Slh1/Cue3 and Slh1/Rqt4 were stably purified as a heterodimer **(Fig. 2c)**, indicating that Cue3 and Rqt4 interact directly with Slh1 independently of each other.

**Fig. 2.**
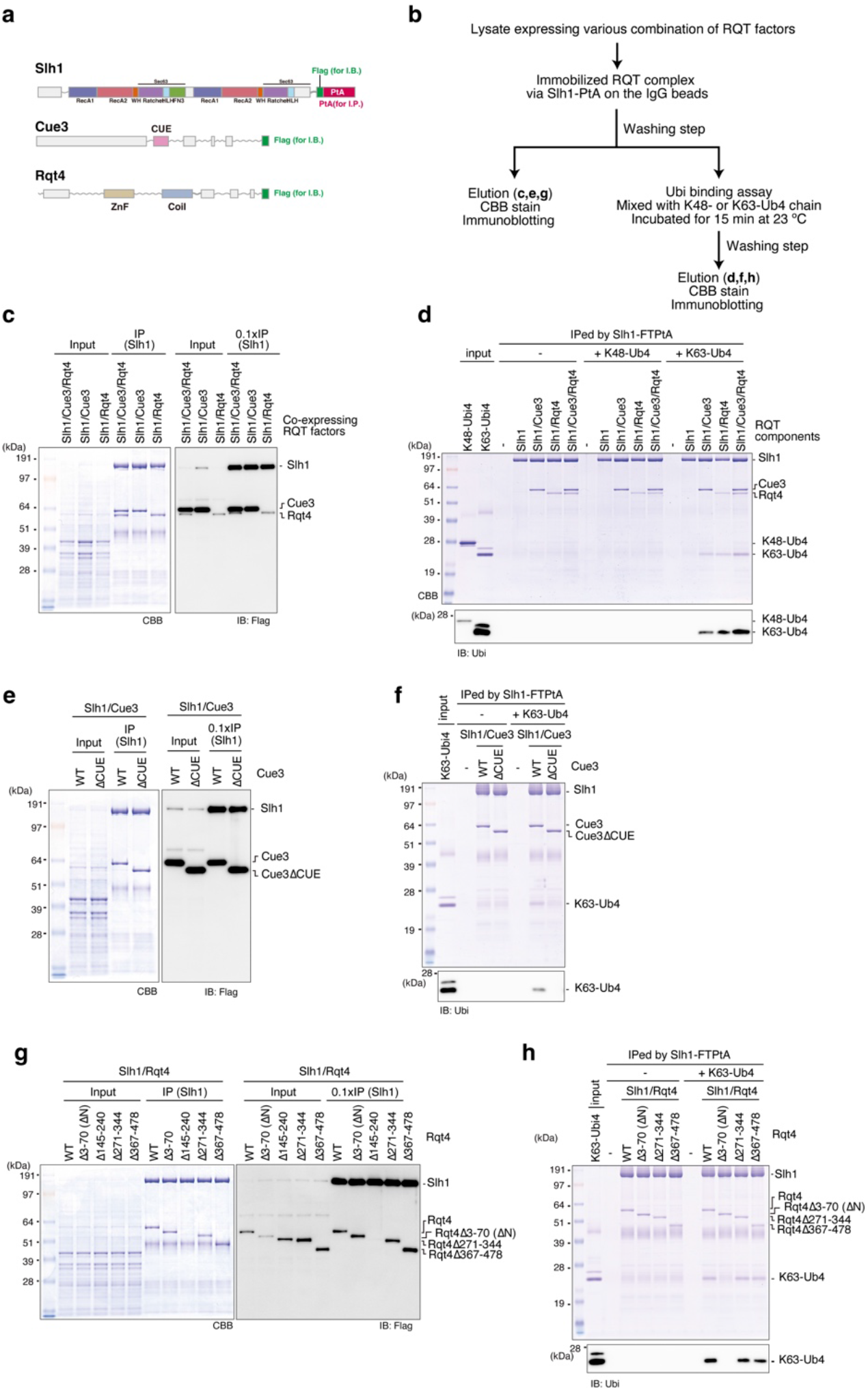
The RQT complex interacts with the K63-linked ubiquitin chain via two accessory proteins. **(a**) Domain structure of each component of the RQT complex. (**b**) Schematic of the experiments. (**c**, **e**, **g**) Purification of the complex containing the indicated RQT factors. Different combinations of the indicated RQT factors were co-expressed in yeast cells. The complex was affinity-purified via Slh1-Flag-TEV-protein A using IgG magnetic beads. Copurified RQT factors were separated by 8% Nu-PAGE and detected by Coomassie brilliant blue (CBB) staining or immunoblotting using an anti-Flag antibody. (**d**, **f**, **h**) Pull-down assay of the RQT complex with the K63- or K48-linked tetraubiquitin chain. The complex containing the indicated RQT factors was immobilized on IgG magnetic beads and mixed with the K63- or K48-linked tetraubiquitin chain. After binding and washing steps (as indicated in 2**b**), the proteins in the final elution were separated by 10% Nu-PAGE and detected by CBB staining or immunoblotting using an anti-ubiquitin antibody.

Next, to test the linkage-specific ubiquitin-binding activity of the RQT complex, Slh1-Flag-TEV-protein A was immobilized on IgG magnetic beads together with various combinations of RQT factors and mixed with K63- or K48-linked tetraubiquitin chains for the pull-down assay **(Fig. 2b)**. Coomassie brilliant blue staining and immunoblotting using an anti-ubiquitin antibody showed that the RQT complex was specifically bound to the K63-linked tetraubiquitin chain but not to the K48-linked tetraubiquitin chain **(Fig. 2d)**. Both the Slh1-Cue3 and the Slh1-Rqt4 heterodimer bound to the K63-linked tetraubiquitin chain **(Fig. 2d)**, whereas Slh1 did not **(Fig. 2d)**. These results indicate that the RQT complex specifically interacted with the K63-linked ubiquitin chain via two accessory proteins, Cue3 and Rqt4.

Because the CUE domain of Cue3 is the ubiquitin-binding domain, we constructed a CUE domain-deleted mutant of Cue3 **(Supplementary Fig. la-b)** and performed a pull-down assay to confirm the interaction between the Slh1-Cue3 complex and the K63-linked ubiquitin chain. Deletion of the CUE domain did not affect the interaction between Cue3 and Slh1 **(Fig. 2e)**; however, its ability to bind to the K63-linked ubiquitin chain was abolished **(Fig. 2f)**. In contrast to Cue3, Rqt4 has no predicted ubiquitin-binding domain and has large IDRs **(Supplementary Fig. le)**. Thus, the domains predicted as the structural regions were selected **(Supplementary Fig. le)** and deleted to construct the Rqt4 mutants **(Supplementary Fig. ld)**. During the purification of different Slh1/Rqt4 mutant complexes, we found that the interaction between Slh1 and Rqt4 was abolished by deletion of the zinc-finger domain located at residues 145-240 of Rqt4 **(Fig. 2g)**. This was consistent with recent cryo-EM data of the RQT complex associated with stalling ribosomes^36^; the zinc-finger domain of Rqt4 directly interacts with the C-terminal RecA1 domain of Slh1 **(Supplementary Fig. 2a, e)**. Because the other deletion mutants retained the ability to form Slh1/Rqt4 complexes **(Fig. 2g)**, we performed pull-down assays to identify the domain mediating the interaction with the K63-linked ubiquitin chain. The results of the pull-down assay showed that the N-terminal domain at residues 3-70 of Rqt4 was essential for interaction with the K63-linked ubiquitin chain **(Fig. 2h)**. Taken together, these results indicate that the RQT complex is equipped with two independent arms, Cue3 and Rqt4, that mediate the interaction with the K63-linked ubiquitin chain.

### The dynamics of the RQT complex

A recent cryo-EM analysis provides structural information of the RQT complex associated with translating ribosomes^36^. However, the flexible regions are not visualized, including the CUE domain of Cue3 and the N-terminal domain of Rqt4. Thus, to understand the dynamics of the RQT complex, we analyzed it using HS-AFM. The largest subunit of the RQT complex, Slh1, belongs to a family of ATP-dependent Ski2-like RNA helicases and contains two RecA-like helicase cassettes **(Fig. 2a and Supplementary Fig. 2a)**, so we first analyzed a single molecule of Slh1. In the HS-AFM images of Slh1, we found two main shapes of the particle, which we termed as Class1 and Class2 particles **(Fig. 3a)**. We next generated the pseudo-AFM images of Class1 and Class2 particles from the Alphafold2 predicted Slh1 structure lacking a flexible N-terminal region **(Fig. 3b)** and compared to actual HS-AFM images of them **(Fig. 3c)**. In Class1 particles, two globular domains, which are consistent with two RecA-like helicase domains, were observed **(Fig. 3b-c and Supplementary Movie 1)**, and the height of pseudo- and actual HS-AFM images were both around 6 nm **(Supplementary Fig. 3a)**. In Class2 particles, the single globular domain seemed to be more tightly contacted with the mica surface, so another globular domain had a higher height **(Fig 3b-c, Supplementary Fig. 3b, and Supplementary Movie 2)**. Collectively, the HS-AFM images of Slh1 showed a structure similar to the pseudo-AFM images **(Fig. 3b-d and Supplementary Fig. 3a-b)**, concluding that the particles in Class1 and Class2 were Slh1. Next, to determine the most major orientation of Slh1 on the mica surface, we classified the Slh1 particles, showing that 65 % and 25 % of the Slh1 particles belong to Class1 and Class2, respectively (**Fig. 3d and Supplementary figure 4**), so, we focused on Classs1 particles as a major particle of Slh1 hereafter.

**Fig. 3.**
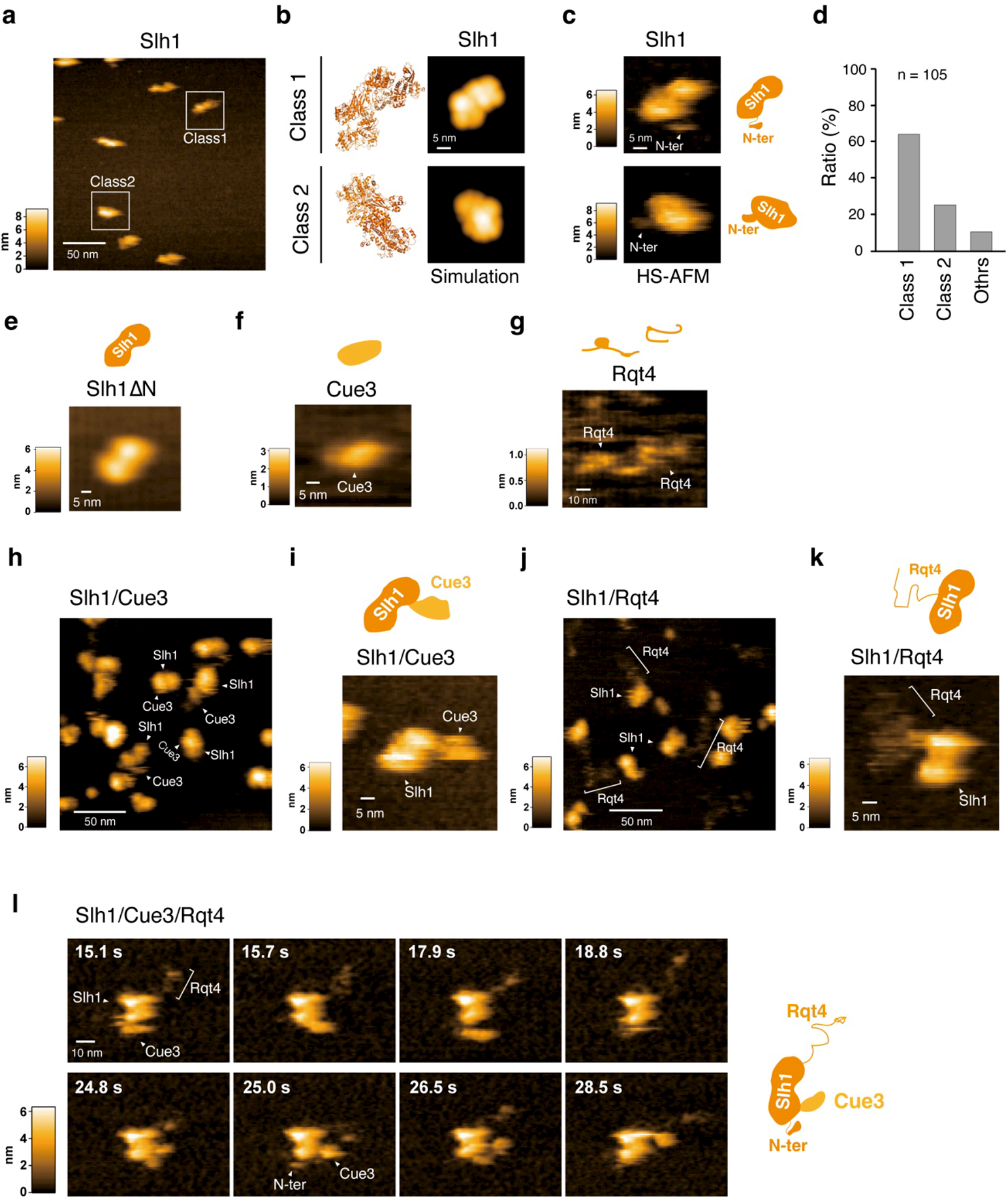
The dynamics of the RQT complex. **(a**) HS-AFM image of Slh1. Two major particles were indicated as Class1 and Class2. **(b)** The pseudo-AFM images of Slh1 belonging to Class1 and Class2 particles, which were simulated using predicted Slh1 structure lacking N-terminal region by Alphafold2. (**c**) The HS-AFM images and schematized molecular features of Slh1. **(d)** Classification of Slh1 particles. (**e**) HS-AFM images of Slh1 lacking N-terminal region (Slh1ΔN). (**f**) HS-AFM image of Cue3. (**g**) HS-AFM image of Rqt4. (**h**, **i**) HS-AFM images of Slh1/Cue3 complex. (**j**, **k**) HS-AFM images of Slh1/Rqt4 complex. (**l**) The time-lapse HS-AFM images of the RQT complex.

**Fig. 4.**
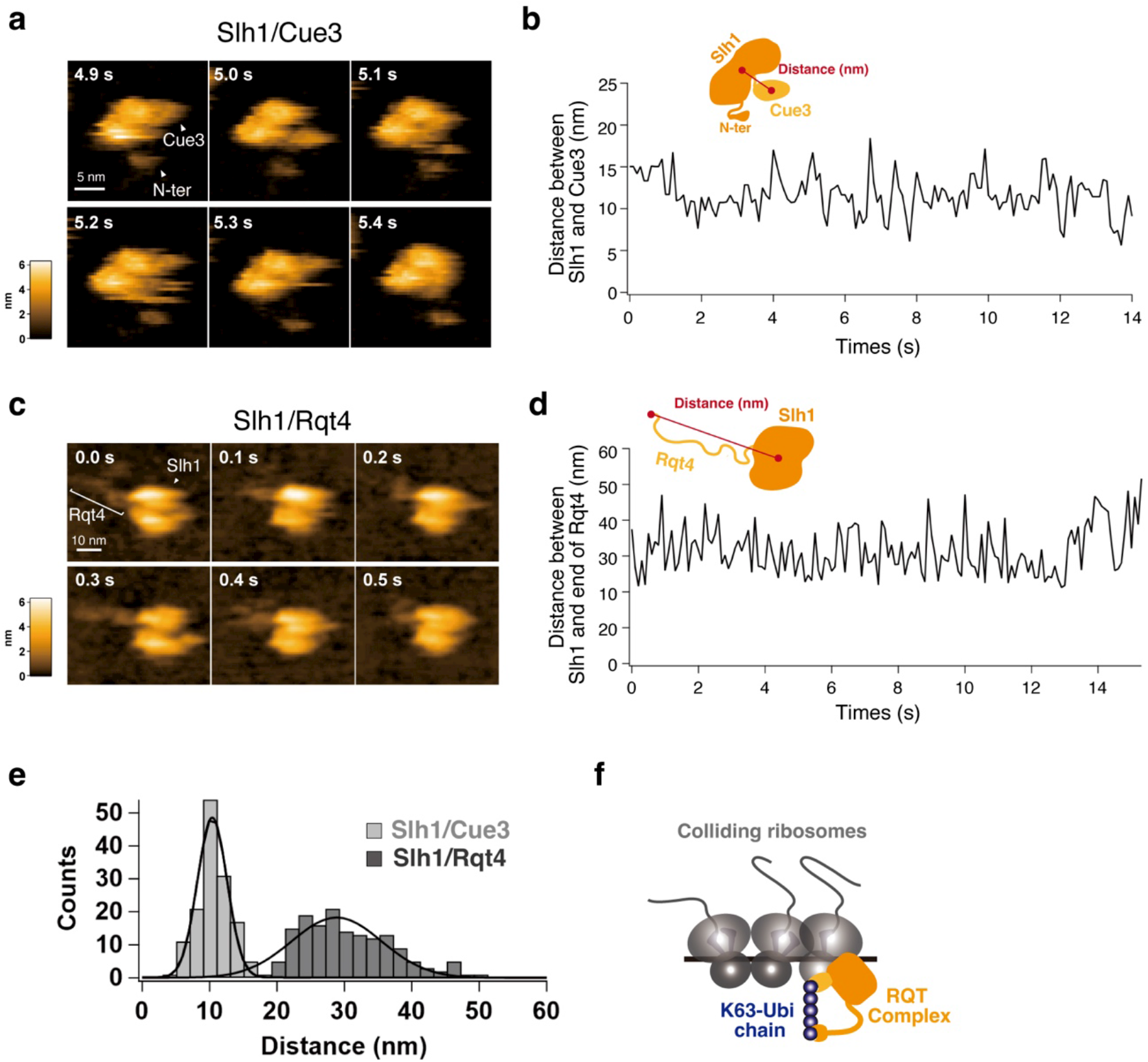
The movable range of the accessory proteins of RQT complex. **(a**) The time-lapse HS-AFM images of the Slh1/Cue3 complex. (**b**) Time course of the distance between Slh1 and Cue3 as described in the schematic diagram. The distance was calculated for each frame per 0.1 second. (**c**) The time-lapse HS-AFM images of the Slh1/Rqt4 complex. (**d**) Time course of the distance between center of Slh1 and the most distant point of Rqt4 from center of Slh1 as described in the schematic diagram. The distance was calculated for each frame per 0.1 second. (**e**) The histogram of the Slh1-Cue3 distance (**b**) and the Slh1-end of Rqt4 distance (**d**).

**Fig. 5.**
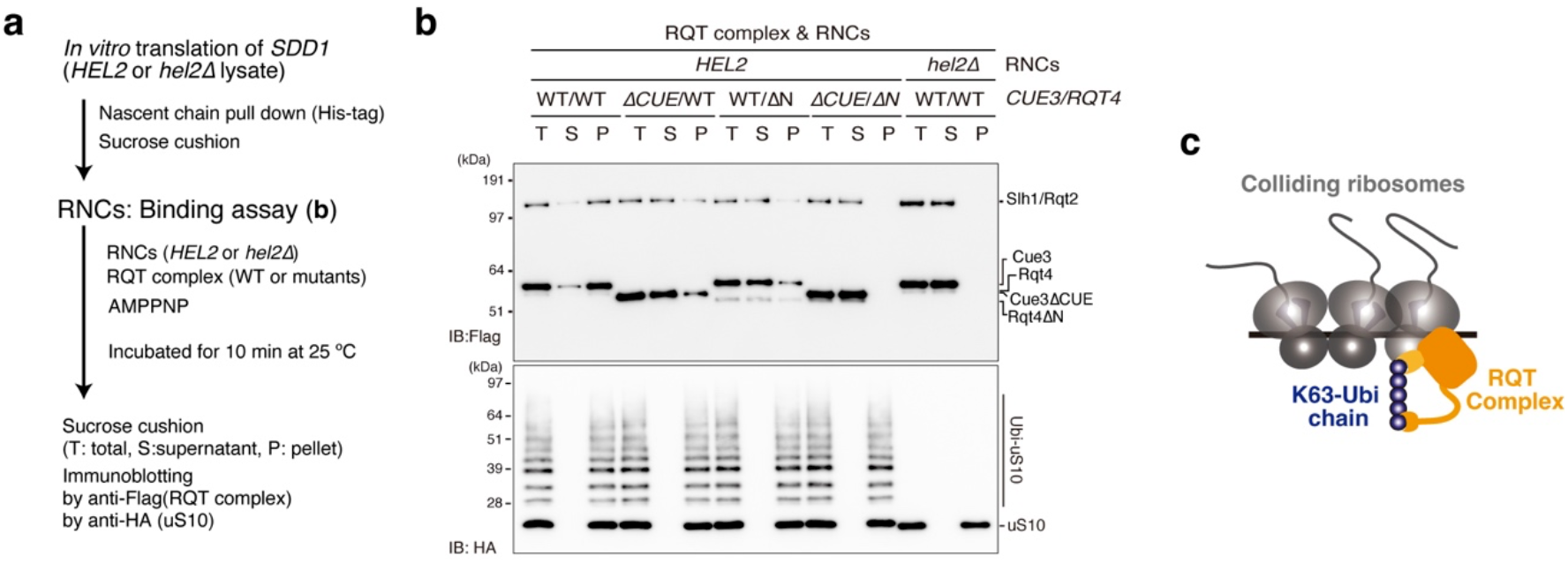
The recruitment of RQT complex into the colliding ribosome in the ubiquitin decoding-dependent manner. **(a)** Schematic of the experiments. **(b)** Binding assay between the colliding ribosome and the RQT complex. The ubiquitinated or nonubiquitinated colliding ribosomes were prepared by *HEL2*-containing or *hel2*-knockout (*hel2*Δ) IVT reaction and mixed with the indicated RQT complex in the presence of AMPPMP as described in 5a. After the binding reaction, free and bound RQT complexes were separated using a sucrose cushion. The RQT factors and uS10 in each fraction were detected by immunoblotting using anti-Flag and anti-HA antibodies, respectively. (**c**) Model of the recognition of colliding ribosomes by the RQT complex.

Although the overall structure of Slh1 seemed stable, a small flexible region was observed **(Fig. 3c and Supplementary Movie 1-2)**. Since the N-terminal of Slh1 is not visualized by cryo-EM analysis^36^, the HS-AFM video might enable visualization of the dynamics of the N-terminal of Slh1. To confirm this possibility, we constructed the Slh1 mutant lacking an N-terminal region (3-217aa: Slh1LN) and analyzed it by HS-AFM. As expected, the fluctuating region completely disappeared in the HS-AFM image and movie of Slh1LN **(Figure 3e and Supplementary Movie 3)**, indicating that the fluctuating region of Slh1 is an N-terminal region.

Analysis of the Slh1/Cue3 heterodimer using HS-AFM suggested that the barrel-shaped protein bound to Slh1 could be Cue3 **(Fig. 3h)**. To confirm this, we focused on the Class1 Slh1 particle and compared the shape of Slh1**(Fig. 3c and Supplementary Movie t)** and Slh1/Cue3 heterodimer **(Fig. 3i and Supplementary Movie 6)**. This comparison clearly showed that the additional globular molecule was associated with Slh1 **(Fig. 3i and Supplementary Movie 6)**. Furthermore, the height of this molecule is around 3 nm, which is consistent with Cue3 **(Fig. 3f, Supplementary Movie 4, and Supplementary Fig. 3c)**, concluding that the associated barrel-shaped molecule is Cue3. The cryo-EM analysis shows that the N-terminal helical domain of Cue3 directly interacts with Slh1 **(Supplementary Fig. 2b)**, whereas the C-terminal domain is not visualized because of flexibility. Consistent with this, the HS-AFM movie of the Slh1/Cue3 complex showed that Cue3 moved around Slh1 **(Supplementary Movie 6)**. Because the disordered probability is high in the C-terminal of Cue3 **(Supplementary Fig. ta)**, we hypothesized that it does not form the structured domain predicted by Alphafold2 **(Supplementary Fig. 2b)**. However, the HS-AFM images and movies of Slh1/Cue3 displayed the flexible but structural Cue3 as a single globular domain in the Slh1/Cue3 complex **(Fig. 3h-i and Supplementary Movie 6)**.

The disorder prediction indicated that Rqt4 contains many IDRs **(Supplementary Fig. 1d)**. Indeed, Rqt4 was visualized as a string protein by the HS-AFM imaging **(Fig. 3g and Supplementary Movie 5)**. The HS-AFM images of the Slh1/Rqt4 complexes also showed a thin-structured accessory protein attached to Slh1, which was believed to be Rqt4 **(Fig. 3j-k)**. In the HS-AFM movie of Slh1/Rqt4 complex, Rqt4 behaved like a super flexible tentacle, which differed from the behavior of Cue3 **(Supplementary Movie 7)**. These observations suggested the assignment of each factor in the Slh1/Cue3/Rqt4 complex **(Fig. 3l)** and provided information on the dynamic properties of the accessory proteins of the RQT complex **(Fig. 3l and Supplementary Movie 8)**.

### The movable range of the accessory proteins of the RQT complex

The dynamic properties of two accessory proteins of the RQT complex seemed to be different **(Fig. 3l and Supplementary Movie 8)**. Cue3 moved around Slh1, but its moving range was limited. By contrast, Rqt4 behaved like a super flexible tentacle; its IDRs seemed to contribute to expanding the searchable areas for ubiquitin detection **(Fig. 3l and Supplementary Movie 8)**. Thus, we next attempted to estimate the searchable range of Cue3 and Rqt4 for the detection of ubiquitin.

To estimate the Cue3 accessible region, we first determined the center position of Slh1 (P1) and Cue3 (P2) using a tracking algorithm, and then the distance between P1 and P2 was calculated for each frame (10 frames per second: fps) as described in Supplementary figure 5. As expected, although Cue3 dynamically moved around Slh1, it stayed near Slh1, which was around 10 nm from the center position of Slh1 **(Fig. 4a-b and Supplementary Movie 6)**. Next, to estimate the Rqt4 accessible region, we defined the center position of Slh1 (P1) and the most distant point of Rqt4 (P3) from P1 and measured the distance between the two points as described in Supplementary figure 6. This revealed that the tentacle of Rqt4 can reach 30 nm away from the center position of Slh1 on average **(Fig. 4c-d and Supplementary Movie 7)**. These observations let us hypothesize that Cue3 interacts with the ubiquitin closest to Slh1 and contributes to engaging the RQT complex at the right position in the ubiquitinated colliding ribosomes and that Rqt4 plays a crucial role in the identification of ubiquitinated colliding ribosomes.

### Ubiquitination is required for recruitment of the RQT complex to the colliding ribosome

According to a simple model, ubiquitination of uS10 is used as a mark for the recruitment of the RQT complex. Given that Rqt4 can compensate for the ubiquitin-binding defect of Cue3, we propose a model by which the ubiquitination of uS10 is recognized by two arms of the RQT complex, which stimulates recruitment of the RQT complex to the colliding ribosome.

To test this hypothesis, ubiquitinated and nonubiquitinated colliding ribosomes were prepared using the *in vitro* translation system, and binding assays between the RQT complex and the colliding ribosomes were performed **(Fig. 5a)**. Affinity-purified RNCs from the *in vitro* translation reaction using *HEL2*-containing (*HEL2*) or *hel2*-knockout (*hel2*Δ) extract were pelleted through a sucrose cushion and then incubated with the RQT complex in the presence of AMPPMP, a nonhydrolyzable ATP analog. To confirm the association of the RQT complex with RNCs, we again pelleted the RNCs through a sucrose cushion and detected the RQT complex in the free- and bound-fraction by immunoblotting. In the ubiquitinated colliding ribosomes (*HEL2*), the wild-type RQT complex was mainly detected in the pelleted fraction **(Fig. 5b)**. By contrast, nonubiquitinated colliding ribosomes prepared from the *hel2*-knockout translation system (*hel2*Δ) did not bind to the RQT complex; almost 100% of the RQT complex remained in the supernatant after sucrose cushion purification **(Fig. 5b)**. Deletion of the CUE domain of Cue3 (Δ*CUE*) or the N-terminal domain of Rqt4 (Δ*N*) decreased binding to the ubiquitinated colliding ribosome **(Fig. 5b)**, whereas double deletion (Δ*CUE*/Δ*N*) abolished binding to the ubiquitinated colliding ribosome **(Fig. 5b)**. Collectively, these results strongly support the model that ubiquitination of uS10 by Hel2 is essential for the recruitment of the RQT complex to the colliding ribosome **(Fig. 5c)**.

### The ubiquitin-binding activity of the RQT complex is essential for the disassembly of colliding ribosomes

To further analyze the role of ubiquitination in the RQC pathway, we monitored the K63-linked ubiquitin-binding activity of the RQT trimer complex lacking the CUE domain of Cue3 or the N-terminal domain of Rqt4. Consistent with the analysis of the heterodimer, a single deletion mutant partially reduced the ubiquitin-binding capacity **(Fig. 6a)**, whereas the double deletion mutant abolished the ubiquitin-binding activity **(Fig. 6a)**. We then performed an *in vivo* RQC reporter assay using the *HA-SDD1-V5* reporter gene^10^. As previously reported ^10^, RQC activity was abolished in the *hel2ΔltnlΔ*, uS10K6/8R*ltnlΔ*, and *slhlΔltnlΔ* mutants **(Fig. 6b)**, and translation arrest products disappeared in these mutants **(Fig. 6b)**. Single deletion of the CUE domain of Cue3 (Δ*CUE*) or the N-terminal domain of Rqt4 (Δ*N*) resulted in the partial reduction of arrest products **(Fig. 6b)**, whereas double deletion (Δ*CUE*/Δ*N*) eliminated the arrest products even in the absence of Ltn1 **(Fig. 6b)**, indicating that the K63-linked ubiquitin signal was independently decoded by the two accessory proteins of the RQT complex, thereby activating the RQC pathway.

**Fig. 6.**
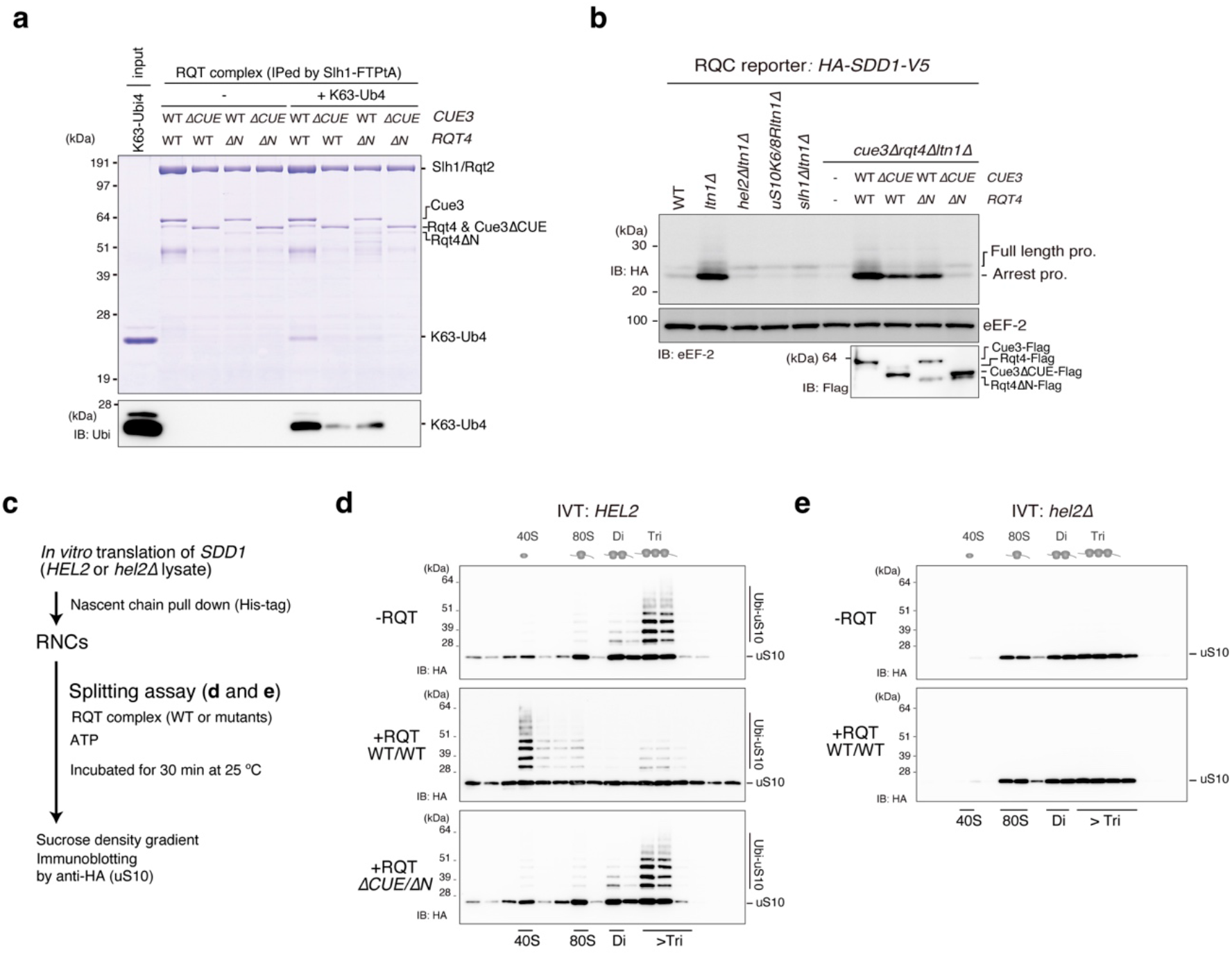
Decoding of K63-linked polyubiquitination is essential for the disassembly of the colliding ribosome by the RQT complex. (**a**) Pull-down assay of the trimer RQT complex with the K63-linked tetraubiquitin chain. The RQT complex was immobilized on IgG magnetic beads and mixed with the K63-linked tetraubiquitin chain. After binding and washing steps, the proteins in the final elution were separated by 10% Nu-PAGE and detected by CBB staining or immunoblotting using an anti-ubiquitin antibody. (**b**) Immunoblotting of the HA-Sdd1-V5 reporter peptide expressed in the deletion strains for RQT- and RQC-related genes. (c) Schematic of the *in vitro* splitting assay. (**d**, **e**) *In vitro* splitting assay using ubiquitinated (*HEL2*) or nonubiquitinated (*hel2Δ*) colliding ribosomes. (**e**). The ubiquitinated or nonubiquitinated colliding ribosomes were prepared by *HEL2*-containing or *hel2*-knockout (*hel2*Δ) IVT reaction and mixed with the indicated RQT complex in the presence of ATP as described in 6c. After the splitting reaction, the ribosomes were separated by sucrose density gradient centrifugation. HA-tagged uS10 in each fraction was detected by immunoblotting using an anti-HA antibody.

To further understand the importance of ubiquitination for the disassembly of the colliding ribosome, we reconstituted the *in vitro* ribosome dissociation assay. First, ubiquitinated and nonubiquitinated colliding ribosomes were prepared from *HEL2*-containing or *hel2*-knockout *in vitro* translation extracts, respectively **(Fig. 6c)**. The purified RNCs were reacted with or without the RQT complex and ATP, and then separated by sucrose density gradient centrifugation **(Fig. 6c)**. As previously reported^10^, the colliding ribosomes forming more than disomes were preferentially ubiquitinated by Hel2 **(Fig. 6d)**. The ubiquitinated colliding ribosomes were disassembled by the RQT complex **(Fig. 6d)**, whereas the nonubiquitinated colliding ribosomes prepared from the *hel2*-knockout *in vitro* translation system were not disassembled **(Fig. 6e)**. Furthermore, the splitting activity of the RQT complex was abolished by double deletion of the CUE domain of Cue3 (Δ*CUE*) and the N-terminal domain of Rqt4 (Δ*N*) **(Fig. 6d)**, indicating that Hel2-mediated K63-linked ubiquitination of uS10 is essential for the recognition and clearance of colliding ribosomes by the RQT complex.

Collectively, we concluded that the RQT complex interacted with the K63-linked ubiquitin chain via two arms composed of Cue3 and Rqt4, and recruited itself into the ubiquitinated colliding ribosome, thereby facilitating the disassembly of the colliding ribosomes using the helicase activity of Slh1 **(Supplementary Fig. 7)**.

## Discussion

The decoding of the ubiquitination in translating ribosomes is the key event in activating the ubiquitin-mediated pathways of translational control. The diverse ubiquitinations of the ribosome are observed; e.g., the ubiquitination of ribosomal proteins uS10, uS3, eS7, and uL23 are essential to initiate different pathways: the RQC pathway^9,11,14^, the 18S non-functional rRNA decay (18S NRD)^37,38^, the no-go decay (NGD) ^18^, and the ribophagy^39^, respectively. However, the decoding mechanism of ribosome ubiquitin-code remains largely unknown. In this study, we focused on the decoding of uS10 ubiquitination in the RQC pathway and revealed that not only Cue3 but also Rqt4 interact with the K63-linked ubiquitin chain and stimulate the recruitment of the RQT complex into the ubiquitinated colliding ribosome. In the previous model, Cue3 is considered as the sole responsible factor for the recognition of uS10 ubiquitination to initiate the RQC pathway, so it is not explained why the deletion of Cue3 does not completely abolish RQC activity^11^. Since Rqt4 contributes to the recognition of the ubiquitinated colliding ribosomes independent of Cue3 recognition, our findings explain the previous discrepancy regarding ubiquitination and its decoder function in the RQC-triggering step.

A recent study demonstrated that ZNF598 functionally marks collided mammalian ribosomes by K63-linked polyubiquitination of uS10 for the trimeric hRQT complex-mediated subunit dissociation^16^, indicating that the crucial role of K63-linked ubiquitination of uS10 for the RQT-mediated clearance is conserved in eukaryotes from yeast to mammals. Since the trimeric hRQT complex is composed of ASCC3 (Slh1), ASCC2 (Cue3), and TRIP4 (Rqt4), the conservation of the mechanism for K63-linked ubiquitin recognition by TRIP4 should be investigated in the future work.

Notably, the visualization of the RQT complex by HS-AFM showed that Rqt4 behaves like a super flexible tentacle, which could search for the K63-linked ubiquitin chain in a wide range of ribosomes. Thus, Rqt4 might recognize the other ubiquitin-codes of the ribosomes. The K63-linked ubiquitin chain is also formed on the uS3 and is essential to inducing the 18S NRD pathway^38^. The ubiquitination of uS3 is initiated by the Mag2-mediated monoubiquitination, followed by K63-linked polyubiquitination by Hel2 or Fap1^37,38^. The deletion of *SLHJ*, which is the core protein of the RQT complex, leads to the reduction of 18S NRD activity^38^, suggesting the involvement of the RQT complex in 18S NRD. Furthermore, the deletion of *RQT4*, but not *CUE3*, partially suppresses the 18S NRD activity^38^, implying that the wide range searchable ability of Rqt4 might enable the recognition of the K63-linked ubiquitin chain of uS3. Future study will uncover the decoding mechanisms of ribosome ubiquitin-code in the 18S NRD.

Recent Cryo-EM analysis provides the RQT complex-associated ribosome structure^36^, which reveals that the RQT complex engages the 40S subunit of the leading ribosome. However, the ubiquitin-binding surface including the CUE domain of Cue3 and N-terminal domain of Rqt4 are not visualized. Here we reported that Cue3 and Rqt4 interact with the K63-linked ubiquitin chain independently of each other, however, it is not yet clear whether Cue3 and Rqt4 bind to the same or different ubiquitin chains on the leading or trailing ribosomes. Thus, the physical interaction between the K63-linked ubiquitin chain of leading or trailing ribosomes and two accessory proteins of the RQT complex is an interesting issue for the future.

## Methods

### Yeast strains and genetic methods

The *S. cerevisiae* strains used in this study are listed in the **Supplementary Table 1**. Gene disruption and C-terminal tagging were performed as previously described^40,4l^.

### Plasmid constructs

All recombinant DNA techniques were performed according to standard procedures using *E coli* DH5α for cloning and plasmid propagation. All cloned DNA fragments generated by PCR amplification were verified by sequencing. Plasmids used in this study are listed in the **Supplementary Table 2**.

### Total protein extraction for the HA-SDD1-V5 reporter assay

Total protein samples for immunoblotting were prepared using the trichloroacetic acid (TCA) precipitation method. For this, exponentially grown yeast cultures were harvested at an OD600 of 0.5-0.8. Cell pellets were resuspended in ice-cold TCA buffer (20 mM Tris pH 8.0, 50 mM NH4OAc, 2 mM EDTA, 1 mM PMSF, and 10% TCA), and then an equal volume of 0.5 mm dia. zirconia/silica beads (BioSpec) was added followed by thorough vortexing for 30 s, three times at 4°C. The supernatant was collected in a new tube. After centrifugation at 18,000 × *g* at 4°C for 10 min and removing the supernatant completely, the pellet was resuspended in TCA sample buffer (120 mM Tris, 3.5% SDS, 14% glycerol, 8 mM EDTA, 120 mM DTT, and 0.01% BPB).

### Immunoblotting

Proteins were separated by SDS-PAGE or Nu-PAGE and transferred to PVDF membranes (Millipore; IPVH00010). After blocking with 5% skim milk, the blots were incubated with the primary antibodies listed in the **Supplementary Table 3**. The secondary antibodies used in this study were conjugated with horseradish peroxidase (HRP) and detected by ImageQuant LAS4000 (Cytiva).

### In vitro translation of SDD1 mRNA

*SDD1 reporter* mRNA was produced using the mMessage mMachine Kit (Thermo Fischer) and used in a yeast cell-free translation extract from *uS10-3HA ski2*Δ and *uS10-3HA ski2*Δ *hel2*Δ cells. This yeast translation extract was prepared, and *in vitro* translation was performed as described previously^42^. The cells were grown in YPD medium to an OD_600_ of 1.5-2.0; washed with water and 1% KCl; and finally incubated with 10 mM DTT in 100 mM Tris, pH 8.0 for 15 min at room temperature. To generate spheroplasts, 2.08 mg zymolyase per 1 g of cell pellet was added in YPD/1 M sorbitol and incubated for 75 min at 30°C. Spheroplasts were then washed three times with YPD/1 M sorbitol and once with 1 M sorbitol, and lysed as described previously^42^ with a douncer in lysis buffer [20 mM HEPES pH 7.5, 100 mM KOAc, 2 mM Mg(OAc)_2_, 10% Glycerol, 1 mM DTT, 0.5 mM PMSF, and complete EDTA-free protease inhibitors (Cytiva)]. From the lysate, an S100 fraction was obtained by low-speed centrifugation followed by ultracentrifugation of the supernatant. The S100 was passed through a PD10 column (Cytiva). *In vitro* translation was performed at 17°C for 60 min using a great excess of template mRNA (20 μg per 200 μl of extract) to prevent degradation of the resulting stalled ribosomes by endogenous response factors.

### Purification of RNCs on the SDD1 mRNA

The stalled RNCs on the *SDD1* mRNA were affinity-purified using the His-tag on the nascent polypeptide chain. After *in vitro* translation at 17°C for 60 min, the extract was applied to Dynabeads™ (Thermo Fischer) for His-tag isolation and pull-down for 5 min at 4°C. The beads were washed with lysis buffer 500 (50 mM HEPES pH 6.8, 500 mM KOAc, 10 mM Mg(OAc)_2_, 0.01% NP-40, and 5 mM B-mercaptoethanol) and eluted in elution buffer (50 mM HEPES pH 7.5, 100 mM KOAc, 2.5 mM Mg(OAc)_2_, 0.01% NP-40, and 5 mM β-mercaptoethanol) containing 300 mM imidazole.

### Sucrose density gradient centrifugation

The purified RNCs were applied to a 15-50% sucrose gradient, and ribosomal fractions were separated by centrifugation for 2 h at 197,568 RCF at 4°C in a SW40 rotor.

### In vitro ubiquitination assay

Ubiquitination reactions were performed in reaction buffer (50 mM Tris-HCl pH 7.5, 100 mM NaCl, 10 mM MgCl_2_, 1 mM DTT, and 50 μM PR619) containing 100 μM ubiquitin (wild-type or mutants; UBPBio), 127 nM Uba1, 500 nM Ubc4, 300 nM Hel2, 30 nM *SDD1*-RNCs, and an energy regenerating source (1 mM ATP, 10 mM creatine phosphate, and 20 μg/ml creatine kinase). Initially, the reaction was mixed at room temperature except for ubiquitin, and ubiquitin was then added and incubated at 28°C for 80 min. To stop the reaction, LDS-sample buffer (Thermo Fischer) was added to the reaction tube. Samples were analyzed by 10% Nu-PAGE and immunoblotting using an anti-HA antibody.

### Purification of Hel2 or Ubal for in vitro ubiquitination assay

Yeast strains overexpressing *Hel2-Flag* or *Uba1-Flag* were cultured in a synthetic complete medium. The harvested cell pellet was frozen in liquid nitrogen and then ground in liquid nitrogen using a mortar. The cell powder was resuspended with lysis buffer 500 (50 mM Tris-HCl pH 7.5, 500 mM NaCl, 10 mM MgCl_2_, 0.01% NP-40, 1 mM DTT, 10 μM ZnCl_2_, and 1 mM PMSF) to prepare the lysate. The lysate was centrifuged at 39,000 g for 30 min at 4°C, and the supernatant fraction was used for the purification step. Hel2-Flag or Uba1-Flag were affinity-purified using anti-DYKDDDDK tag antibody beads (WAKO). After the washing step with stepwise concentration of NaCl from 500 mM to 100 mM, Hel2-FLAG was eluted from the beads using elution buffer containing 250 μg/ml Flag peptides at 4°C for 1 h.

### Purification of Ubc4 for the in vitro ubiquitination assay

Recombinant Ubc4 was purified as GST-Ubc4 from *E. coli* Rossetta-gami 2 (DE3) harboring pGEX-Ubc4. The cell pellet was resuspended with lysis buffer 500 and disrupted by sonication to prepare the lysate. The lysate was centrifuged at 39,000 g for 30 min at 4°C, and the supernatant fraction was used for the purification step. GST-Ubc4 was affinity-purified using Glutathione Sepharose 4B (Cytiva). After the washing step with a stepwise concentration of NaCl from 500 mM to 100 mM, no tagged Ubc4 was eluted by PreScission Protease (Cytiva).

### Pull-down assay of the RQT complex

The yeast strain overexpressing various combinations of RQT factors or their mutants (*SLH1-FTP*, *CUE3-Flag*, *RQT4-Flag*) were cultured in synthetic complete medium. The harvested cell pellet was frozen in liquid nitrogen and then ground in liquid nitrogen using a mortar. The cell powder was resuspended with RQT-R buffer (50 mM Tris pH7.5, 100 mM NaCl, 2.5 mM MgCl_2_, 0.1% NP-40, 10% glycerol, 100 mM L-arginine, 1 mM DTT, 10 μM ZnCl_2_, and 1 mM PMSF) to prepare the lysate. The lysate was centrifuged at 39,000 g for 30 min at 4°C, and the supernatant fraction was used for the purification step. The RQT complex was affinity-purified using IgG beads (Cytiva), followed by elution using LDS-sample buffer (Thermo Fischer). The elution was separated by 8% Nu-PAGE and analyzed by CBB staining or immunoblotting using an anti-Flag antibody (Sigma).

### Pull-down assay of the RQT complex with the tetra-ubiquitin chain

The RQT complex was immobilized with IgG beads (Cytiva) as described in the section of the pull-down assay of the RQT complex. The immobilized RQT complex was mixed with K48- or K63-linked tetraubiquitin (UBPBio) in RQT-R buffer and incubated for 15 min at 23°C. After the reaction, the immobilized RQT complex with IgG beads was washed with RQT-R buffer and eluted with LDS-sample buffer (Thermo Fischer). The elution was separated by 10% Nu-PAGE and analyzed by CBB staining or immunoblotting using an antiubiquitin antibody (Santa Cruz Bio.).

### HS-AFM imaging and analysis

A freshly cleaved mica surface was prepared by removing the top layers of mica using Scotch tape. A drop of the RQT complex (2 μl: 5 nM) in RQT-AFM buffer (50 mM Tris pH 7.5, 100 mM NaCl, 10 mM MgCl_2_, 100 mM L-arginine, and 10 μM ZnCl_2_) was deposited onto the mica surface. After incubation for 5 min, the mica surface was rinsed twice with 20 μl RQT-AFM buffer, and then the sample stage was immersed in a liquid cell containing 130 μl RQT-AFM buffer. HS-AFM imaging was performed at room temperature on an MS-NEX (RIBM) using two different types of short cantilevers (USF-F1.2-k0.15-10, NanoWorld: a spring constant ∼0.15 N/m, a resonance frequency ~ 0.6 MHz, AC-7, Olympus: a spring constant ~0.1Nim, a resonance frequency ~ 0.6). The HS-AFM data were acquired using the Eagle software package by RIBM in an Igor Pro-6 (WeveMetrics).

HS-AFM images were viewed and analyzed using the laboratory-made analysis software based on Igor Pro-9 (WaveMetrics). To estimate the searchable area of Cue3, firstly the center positions of Slh1 (P1) and Cue3 (P2) in the first frame were calculated from the center of gravity, then each center position was independently determined for each frame using a tracking algorithm based on 2D correlation coefficients, and the distance between P1 and P2 was calculated for each frame (**Supplementary Fig. 5**). To estimate the searchable area of Rqt4, first, the center position (P1) of Slh1 in each frame was determined using the same way described above. Next, the outline of the Slh1/Rqt4 was detected with an manually adjusted threshold for each frame and calculated the distance beteem the center position (P1) of Slh1 and the most distant point P3 corresponding to a portion of Rqt4 within the outline (**Supplementary Fig. 6**).

### In vitro binding assay between SDDI-RNCs and the RQT complex

The pure *SDD1*-RNCs pelleted through the sucrose cushion were reacted with the RQT complex including various combinations of RQT factors together with 1 mM AMP-PNP. After incubation for 10 min at 25°C, the *SDD1*-RNCs were again pelleted through a sucrose cushion. The *SDD1*-RNCs and RQT complex were detected by immunoblotting using anti-HA and anti-Flag antibodies.

### In vitro splitting assay

The purified RNCs (100 nM ribosome) containing 100 nM Rqc2 were incubated in various combinations with 10 nM RQT complex and 1 mM ATP in reaction buffer (50 mM HEPES, pH 7.4, 100 mM KOAc, 2.5 mM Mg(OAc)_2_, 0.01% NP-40, 5 mM β-mercaptoethanol, and 0.2 Uiμl SUPERase In RNase inhibitor) for 45 min at 25°C. After incubation, ribosomal fractions were separated by sucrose density gradient centrifugation. The RNCs were monitored by measuring UV absorbance at 254 nm, and the ubiquitinated uS10 in each fraction was detected by immunoblotting using an anti-HA antibody.

### Purification of Rqc2 for the in vitro splitting assay

The yeast strain overexpressing *RQC2-FTP* was cultured in synthetic complete medium. The harvested cell pellet was frozen in liquid nitrogen and then ground in liquid nitrogen using a mortar. The cell powder was resuspended with lysis buffer 500 to prepare the lysate. The lysate was centrifuged at 39,000 g for 30 min at 4°C, and the supernatant fraction was used for the purification step. Rqc2-FTP was affinity-purified using IgG beads (Cytiva), followed by TEV protease cleavage to release it, and then repurified by anti-DYKDDDDK tag antibody beads (WAKO).

### Purification of the RQT complex for the in vitro splitting assay and HS-AFM

Yeast strains overexpressing wild-type RQT complex (*SLH1-FTP*, *CUE3*, *RQT4*) or mutant RQT complex (*SLH1-FTP*, *CUE3ΔCUE*, *RQT4ΔN*) were cultured in synthetic complete medium. The harvested cell pellet was frozen in liquid nitrogen and then ground in liquid nitrogen using a mortar. The cell powder was resuspended with RQT buffer (50 mM HEPES pH 7.4, 00 mM KOAc, 2.5 mM MgOAc, 0. NP-40, 0 glycerol, 00 mM -arginine, 5 mM DTT, 0 M nCl, and mM PMSF) to prepare the lysate. The lysate was centrifuged at 39,000 g for 30 min at 4°C, and the supernatant fraction was used for the purification step. The RQT complex was affinity-purified using IgG beads (Cytiva), followed by TEV protease cleavage to release it. The TEV elution was further incubated with benzonase (MERCK) to digest the ribosome and then repurified by anti-DYKDDDDK tag antibody beads (WAKO).

## Data Availability

Any data relating to the findings presented in this Article are available within the article and its supplementary information files. Additional information and relevant raw data are available from the corresponding author upon reasonable request. Source data are provided with this paper.

## Code Availability

The code used to analyze the data in Figure 4 are available from the corresponding author upon reasonable request.

## Acknowledgments

We thank Dr. Kodera N. and Dr. Tmai T. for the excellent technical assistance regarding the use of HS-AFM in the 8^th^ Bio-AFM summer school at Kanazawa University. This study was supported by JST PREST Grant Number JPMJPR21EE to Y.M. and by MEXT/JSPS KAKENHT Grant Number 21H05710, 21H00267, 22H02606 to Y.M., JP21H01772, JP21H00393 to T. U. This work was supported by AMED Grant Number JP 20gm1110010h0002 (T.I.), MEXT/JSPS KAKENHI Grant Numbers JP19H05281, 21H05277, 22H00401 (T.I.), Research grants from Takeda Science Foundation (T.I.).

## Author contributions

The biochemical and genetic experiments were designed and performed by Y.M.; the analyses of HS-AFM were performed by Y.M. and T.U.; the algorithms used for the analysis of HS-AFM images were built by T.U.; the results were interpreted by Y.M., T.U., and T.T.; Y.M. and T.T. wrote the manuscript.; and T.T. supervised the project.

