## Supplementary information for "Decoding of the ubiquitin code for clearance of colliding ribosomes by the RQT complex"

**This PDF file includes:**

Supplementary Figs 1 to 7

Supplementary Table 1 to 3

Supplementary Movie 1 to 8

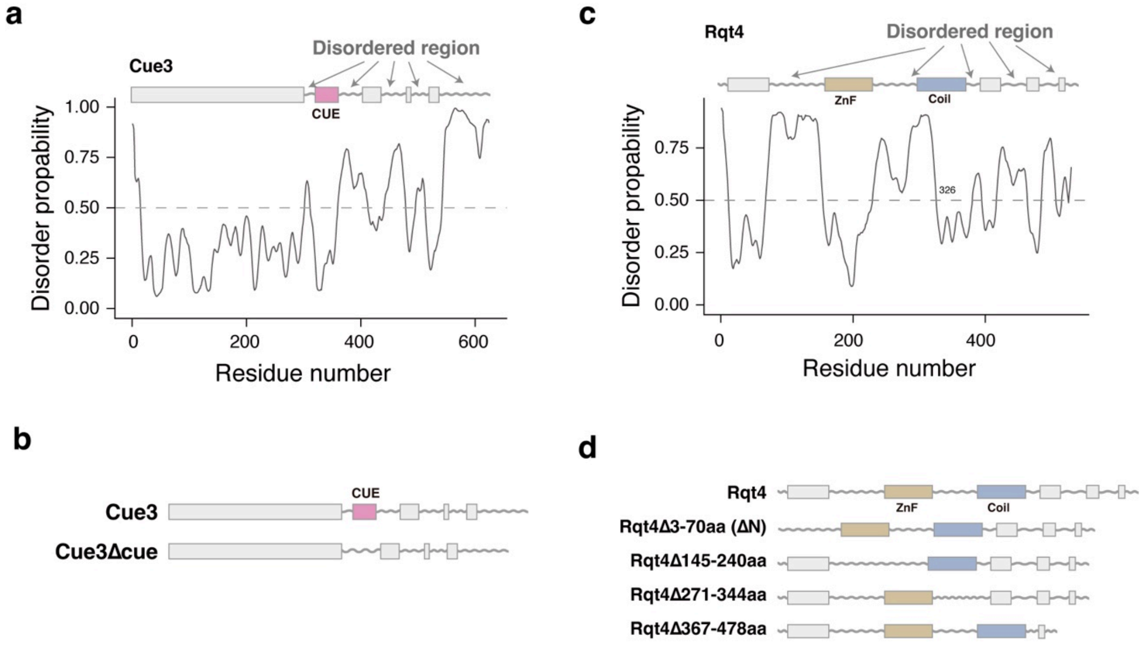


**Supplementary Figure 1. Domain structure of two accessory proteins in the RQT complex**

**(a)** Domain diagram of Cue3 and the order/disorder map along its length as predicted by PrDOS (https://prdos.hgc.jp/cgi-bin/top.cgi) [43]. **(b)** Domain diagram of Cue3 and its mutants. **(c)** Domain diagram of Rqt4 and the order/disorder map along its length as predicted by PrDOS[43]. **(d)** Domain diagram of Rqt4 and its mutants.

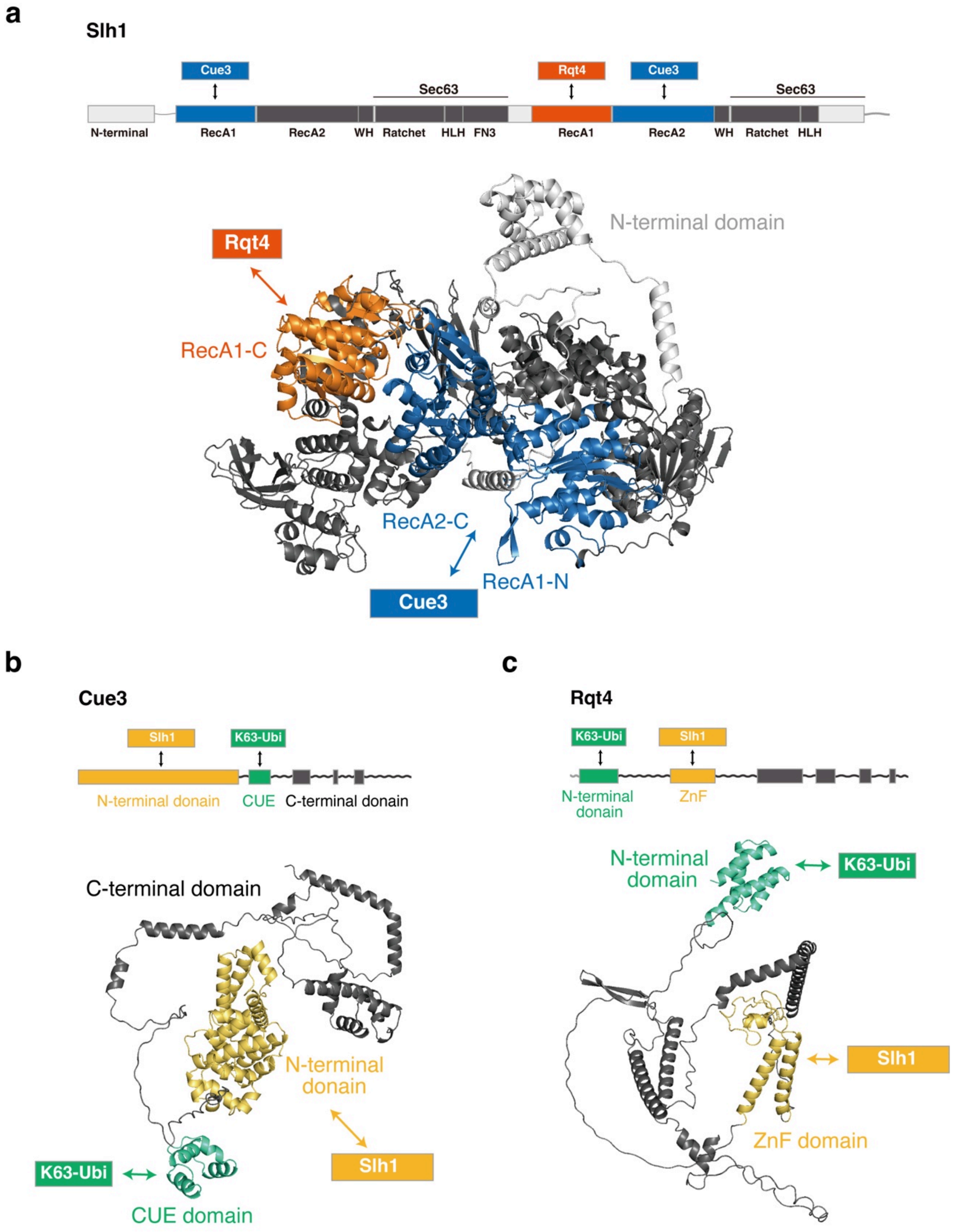


**Supplementary Figure 2. Structure model of RQT components predicted by AlphaFold2**

Domain diagram and 3D structure model of Slh1 **(a)**, Cue3 **(b)**, and Rqt4 **(c)** were predicted by Alphafold2[44,45]. The interacting domains among RQT factors are indicated[34].

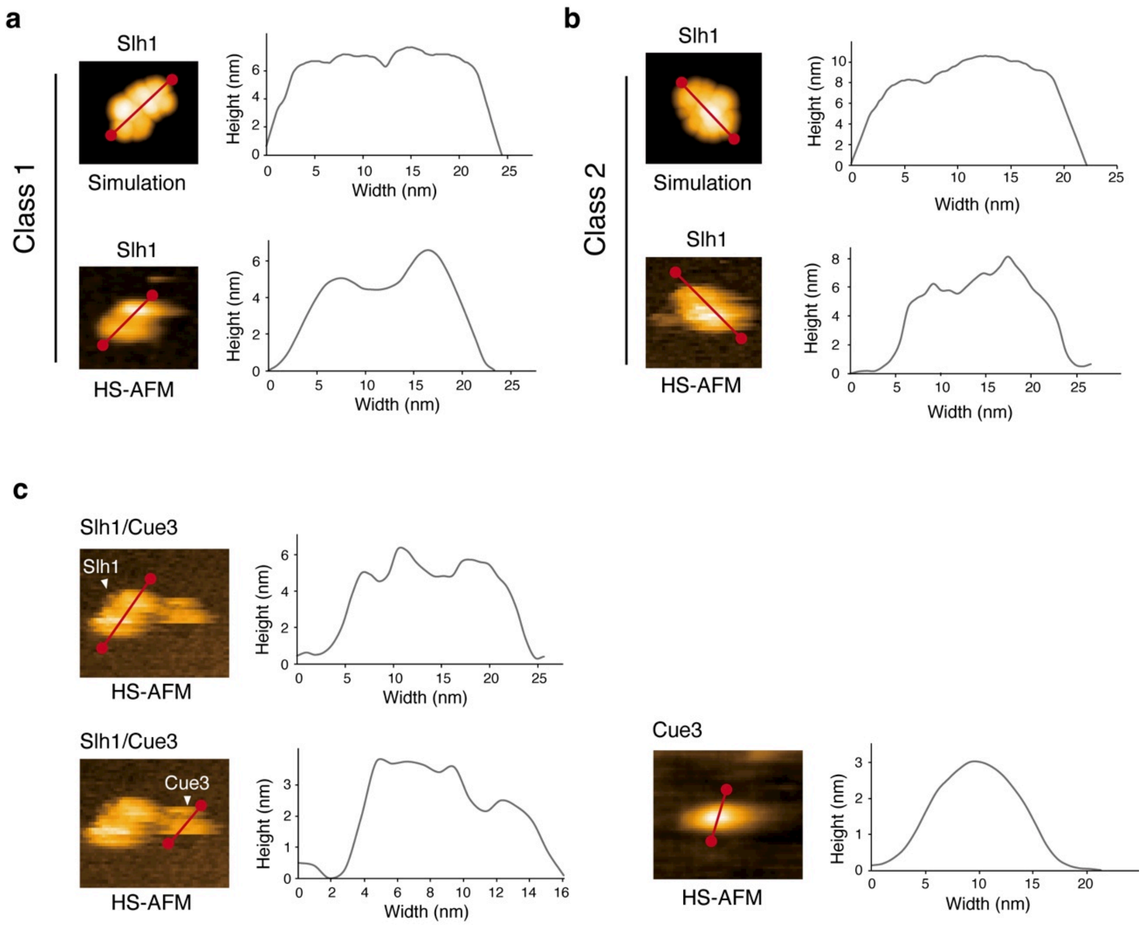


**Supplementary Figure 3. Height analysis of HS-AFM images of Slh1 and Slh1/Cue3 complex**

(**a**) Pseudo- and actual HS-AFM images of Slh1 belonging to Class1 particle. Line graph showing a height profile along the red line on pseudo- and actual HS-AFM images of Slh1 Class1 particle. (**b**) Pseudo- and actual HS-AFM images of Slh1 belonging to Class2 particle. Line graph showing a height profile along the red line on pseudo- and actual HS-AFM images of Slh1 Class2 particle. (**c**) HS-AFM images of Slh1/Cue3 and Cue3. Line graph showing a height profile along the red line on p HS-AFM images of Slh1/Cue3 complex and Cue3.

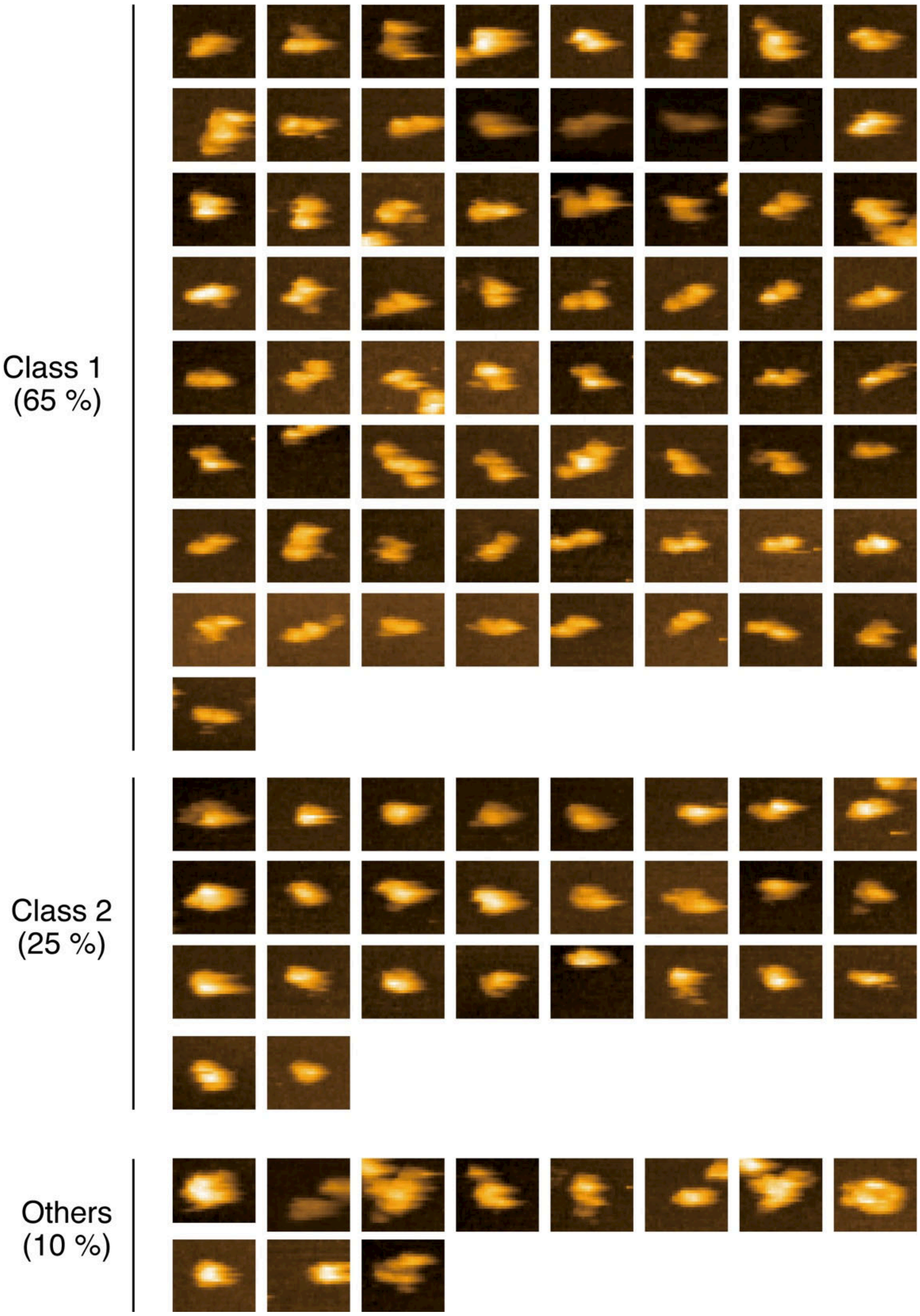


**Supplementary Figure 4. HS-AFM images of Slh1 particle for Classification**

The classification results in shown in Fig. 3d

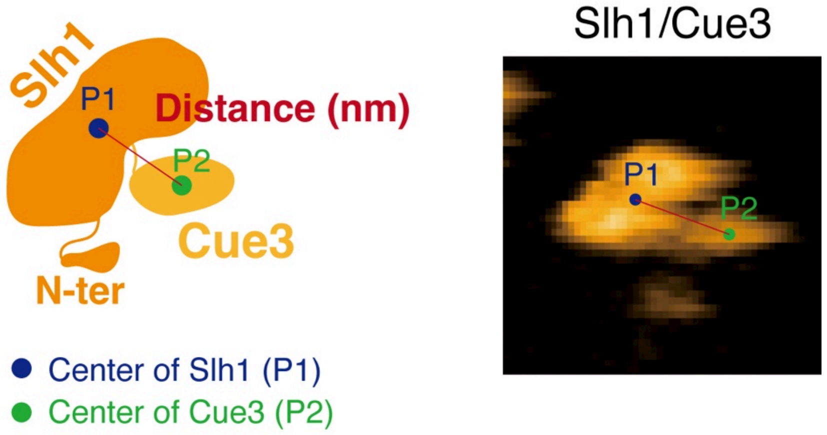


**Supplementary Figure 5. The method used to obtain the distance between Slh1 and Cue3.**
The center positions of Slh1 (P1) and Cue3 (P2) are determined by a tracking algorithm, and then the distance between P1 and P2 is calculated for each frame.

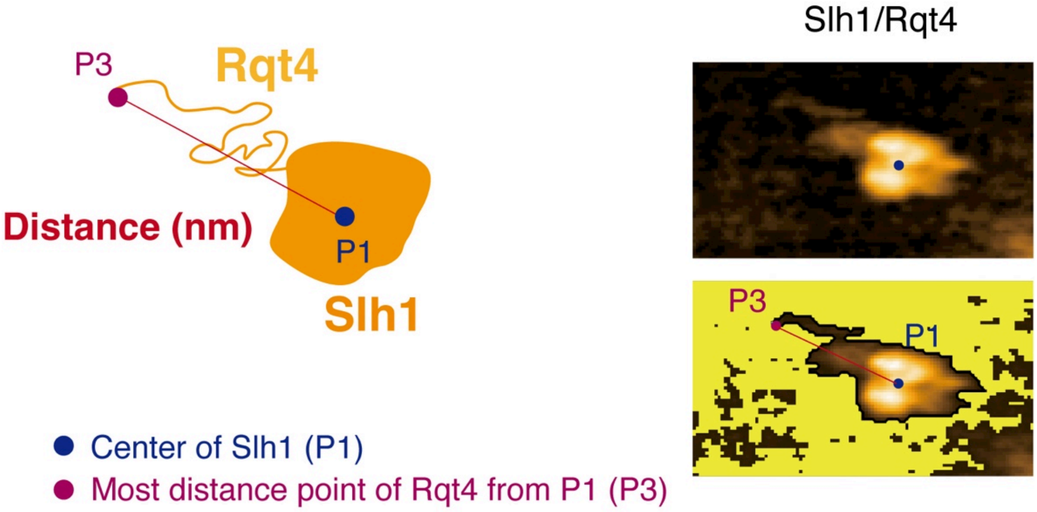


**Supplementary Figure 6. The method used to obtain the distance between center of Slh1 and the most distant point of Rqt4.**

The center position of Slh1 (P1) is determined by a tracking algorithm. To visualize the Rqt4, we manually set the threshold to remove the background and identify the region of Rqt4 as shown by the yellow-field region. The most distant point of Rqt4 from P1 (P3) was determined using an algorithm, and then the distance between P1 and P3 was calculated for each frame.

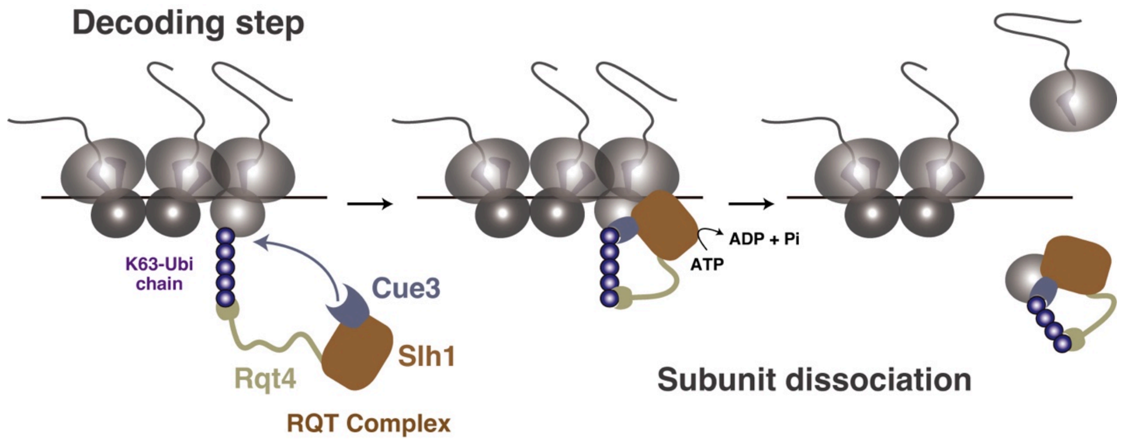


**Supplementary Figure 7. Model of the decoding step of the K63-linked polyubiquitinated colliding ribosome by the RQT complex**

**Supplementary Table 1: The yeast strains used in this study**

| Name | Genotype | Source |
|---|---|---|
| W303a | *MATa ade2 his3 leu2 trp1 ura3 can1* | Lab.stock |
| *ski2ΔuS10-3HA* | *W303a ski2Δ::kanMX4 uS10-3HA::HISMX6* | Ref.10 |
| *ski2ΔuS10-3HAhel2Δ* | *W303a ski2Δ::kanMX4 uS10-3HA::HISMX6 hel2Δ::natMX4* | This study |
| *ltn1Δ* | *W303a ltn1Δ::kanMX4* | Ref.11 |
| *ltn1Δhel2Δ* | *W303a hel2Δ::natMX4 ltn1Δ::kanMX4* | Ref.11 |
| *ltn1ΔuS10 shuffle* | *W303a us10Δ::natMX4 ltn1Δ::kanMX4* | Ref.11 |
| *ltn1Δslh1Δ* | *W303a slh1Δ::natMX4 ltn1Δ::kanMX4* | Ref.11 |
| *ltn1Δcue3Δrqt4Δ* | *W303a cue3::kanMX4 rqt4Δ::natMX4 ltn1Δ::hygMX4* | Ref.11 |

**Supplementary Table 2: The plasmids used in this study**

| Name | Feature | Source |
|---|---|---|
| p415-Hel2-Flag | CEN, LEU2, GPD promoter, HEL2-FLAG | Ref.18 |
| pGEX-Ubc4 | pGEX, AmpR, tac promoter, GST-3C-UBC4 | Ref.18 |
| p416-Uba1-Flag | CEN, URA3, GPD promoter, UBA1-FLAG | This study |
| p425GAL-Slh1-FTP | 2μ, LEU2, GAL1 promoter, Slh1-FTP | Ref.10 |
| p425GAL-Slh1ΔN-FTP | 2μ, LEU2, GAL1 promoter, Slh1ΔN-FTP | This study |
| p424GAL-Cue3 | 2μ, TRP1, GAL1 promoter, Cue3 | Ref.10 |
| p424GAL-Cue3-Flag | 2μ, TRP1, GAL1 promoter, Cue3-Flag | This study |
| p424GAL-Cue3Δcue-Flag | 2μ, TRP1, GAL1 promoter, Cue3ΔCUE-Flag | This study |
| p426GAL-Rqt4 | 2μ, URA3, GAL1 promoter, Rqt4 | Ref.10 |
| p426GAL-Rqt4-Flag | 2μ, URA3, GAL1 promoter, Rqt4-Flag | This study |
| p426GAL-Rqt4Δ70(ΔN)-Flag | 2μ, URA3, GAL1 promoter, Rqt4Δ70aa-Flag | This study |
| p426GAL-Rqt4Δ145-240-Flag | 2μ, URA3, GAL1 promoter, Rqt4Δ145-240aa-Flag | This study |
| p426GAL-Rqt4Δ271-344-Flag | 2μ, URA3, GAL1 promoter, Rqt4-Δ271-344aaFlag | This study |
| p426GAL-Rqt4Δ367-478-Flag | 2μ, URA3, GAL1 promoter, Rqt4Δ367-478aa-Flag | This study |
| pRS415-Rqt4-Flag | CEN, LEU2, Rqt4 promoter, Rqt4-Flag | Ref.11 |
| pRS415-Rqt4Δ70(ΔN)-Flag | CEN, LEU2, Rqt4 promoter, Rqt4Δ70aa-Flag | This study |
| pRS414-Cue3-Flag | CEN, TRP1, Cue3 promoter, Cue3-Flag | Ref.11 |
| pRS414-Cue3ΔCUE-Flag | CEN, TRP1, Cue3 promoter, Cue3ΔCUE-Flag | Ref.11 |
| p416GPD-HA-SDD1-V5 | CEN, URA3, GPD promoter, HA-SDD1-V5 | Ref.10 |

**Supplementary Table 3: The antibodies used in this study**

| Antibody | Company | RRID | Dilution |
|---|---|---|---|
| Anti-HA antibody | Roche | RRID: AB_390917 | 1:10000 |
| Anti-Flag antibody | Sigma-Aldrich | RRID: AB_262044 | 1:5000 |
| Anti-ubiquitin antibody | Santa Cruz Biotechnology | RRID: AB_628423 | 1:1000 |
| Antiubiquitin (linkage-specific K48) antibody | Abcam | RRID: AB_2783797 | 1:1000 |
| Antiubiquitin (linkage-specific K63) antibody | Abcam | RRID: AB_2895239 | 1:1000 |
| Anti-eEF-2 antibody | Lab.stock | N/A | 1:20000 |

**Supplementary Movie 1: HS-AFM movie of Slh1 Class1 particle**

**Supplementary Movie 2: HS-AFM movie of Slh1 Class2 particle**

**Supplementary Movie 3: HS-AFM movie of Slh1ΔN**

**Supplementary Movie 4: HS-AFM movie of Cue3**

**Supplementary Movie 5: HS-AFM movie of Rq4**

**Supplementary Movie 6: HS-AFM movie of Slh1/Cue3 complex**

**Supplementary Movie 7: HS-AFM movie of Slh1/Rqt4 complex**

**Supplementary Movie 8: HS-AFM movie of Slh1/Cue3/Rqt4 complex**
